# A molecularly engineered lectin destroys EGFR and inhibits the growth of non-small cell lung cancer

**DOI:** 10.1101/2024.03.18.585535

**Authors:** Susana M. Chan, Zoe Raglow, Anupama Pal, Scott D. Gitlin, Maureen Legendre, Dafydd Thomas, Ranjit K Mehta, Mingjia Tan, Mukesh K. Nyati, Alnawaz Rehemtulla, David M. Markovitz

## Abstract

Survival rates for non-small cell lung cancer (NSCLC) remain low despite the advent of novel therapeutics. Tyrosine kinase inhibitors (TKIs) targeting mutant epidermal growth factor receptor (EGFR) in NSCLC have significantly improved mortality but are plagued with challenges--they can only be used in the small fraction of patients who have susceptible driver mutations, and resistance inevitably develops.

Aberrant glycosylation on the surface of cancer cells is an attractive therapeutic target as these abnormal glycosylation patterns are typically specific to cancer cells and are not present on healthy cells. H84T BanLec (H84T), a lectin previously engineered by our group to separate its antiviral activity from its mitogenicity, exhibits precision binding of high mannose, an abnormal glycan present on the surface of many cancer cells, including NSCLC.

Here, we show that H84T binds to and inhibits the growth of diverse NSCLC cell lines by inducing lysosomal degradation of EGFR and leading to cancer cell death through autophagy. This is a mechanism distinct from EGFR TKIs and is independent of EGFR mutation status; H84T inhibited proliferation of both cell lines expressing wild type EGFR and those expressing mutant EGFR that is resistant to all TKIs. Further, H84T binds strongly to multiple and diverse clinical samples of both pulmonary adenocarcinoma and squamous cell carcinoma. H84T is thus a promising potential therapeutic in NSCLC, with the ability to circumvent the challenges currently faced by EGFR TKIs.

## Introduction

Lung cancer is the leading cause of cancer-related death worldwide, and non-small cell lung cancer (NSCLC) accounts for more than 80% of lung cancer diagnoses (1). Though the treatment of NSCLC has been revolutionized by the development of targeted molecular therapies and immune checkpoint inhibitors, 5-year survival rates remain less than 20% (2).

Epidermal growth factor receptor (EGFR), a member of the HER/erbB family of receptor tyrosine kinases, is overexpressed in a variety of epithelial cancers, including NSCLC (3). Ligand binding activates the tyrosine kinase, which then triggers downstream signaling pathways (including RAS-RAF-MEK-MAPK, STAT, and PI3K-PTEN-AKT), and ultimately leads to increased cell proliferation, survival, and metastasis (4). Tyrosine kinase inhibitors (TKIs) targeting mutant EGFR in NSCLC have undergone significant evolution since their introduction in the early 2000s (5, 6). The third generation EGFR TKI osimertinib is now first-line therapy for patients with EGFR-mutant NSCLC and has significantly prolonged progression-free and overall survival, both versus standard chemotherapy and first-generation TKIs (7, 8). However, overall survival remains low at 38 months even with osimertinib (9). More critically, development of resistance to TKIs, including osimertinib, is inevitable, and usually occurs about 10 months after initiation of therapy. Resistance is acquired through multiple mechanisms, including development of mutations in the osimertinib binding site and involvement of more complex off-target resistance pathways, but not all resistance mechanisms are well understood (10). Additionally, TKIs targeting EGFR are only effective in a minority of NSCLC patients; EGFR is commonly overexpressed in NSCLC, but the driver mutations necessary to confer susceptibility to these agents occur in only 10-15% of Caucasian patients (versus around 30-60% of Asian patients) (11). Thus, while targeted EGFR therapies have provided much needed advancement in the treatment of NSCLC, these treatments face significant challenges, and new therapeutic approaches are urgently needed.

Glycosylation, the process by which sugars are enzymatically linked to proteins, lipids, and RNA, is highly dysregulated in cancer and is involved in mediating many aspects of oncogenesis (12). Aberrant glycosylation on the surface of cancer cells is a promising therapeutic target, as expression of certain carbohydrate structures is usually specific to tumor cells and not found on normal tissue. Indeed, novel cancer therapeutics targeting abnormal glycosylation patterns have demonstrated efficacy in a variety of cancer types (12). EGFR is known to exhibit anomalous glycosylation patterns in tumor cells, including aberrant expression of high mannose (13, 14). High mannose is a type N-glycan consisting of 5-9 mannose molecules. In normal cells, high mannose is present early in glycan synthesis, but is clipped off in the Golgi apparatus; this clipping is often absent in cancer cells, leaving high mannose present on cancer (and stromal) cell surfaces (12). High mannose is known to be present on NSCLC-derived EGFR, and has already proved to be a potentially attractive therapeutic target (13).

A significant issue in therapeutic glycan-targeting is the specific recognition and binding of aberrant glycans by the therapeutic agent. While antibodies (or antibody fragments) have historically been used, they have a number of limitations, including lack of sufficient specificity for the target glycan (15, 16). Lectins are proteins that have evolved over millions of years to bind glycans with high specificity and are therefore uniquely poised to target anomalous glycans on cancer cells (12). In the past, therapeutic utility of lectins has been limited by their mitogenicity. However, we previously demonstrated that two essential properties of a lectin – mitogenicity and therapeutic (in this first case antiviral) activity – can be separated through targeted molecular engineering. This approach led to the creation of H84T BanLec (H84T), a non-mitogenic lectin derived from a mitogenic, naturally-occurring banana lectin (BanLec), which exhibits potent antiviral activity (17–22). The separation of these activities was achieved by mutating a histidine residue at position 84 to threonine (hence H84T). H84T exerts its antiviral activity by binding high mannose present on a variety of viral surfaces (19, 20). H84T is well tolerated in mouse models, exerts tissue specificity for the lung when administered systemically, has minimal off-target effects, and has a long half-life (17, 18, 23).

Given this specificity for high mannose, which is also present in a variety of cancers including NSCLC, and favorable therapeutic profile in mouse models, we hypothesized that H84T could be an effective therapy for NSCLC, perhaps by binding to the cancer-specific high-mannose covering EGFR and inactivating the tyrosine kinase activity.

## Experimental Procedures

### Cell lines and cell culture

The NSCLC cell lines A549, NCI-H1975 (H1975) and HCC827 were cultured in RPMI (Thermo Fisher Scientific (TFS), catalog #11875-093), supplemented with 10% fetal bovine serum (FBS) and 1X penicillin-streptomycin, at 37°C in a 5% CO_2_ cell culture incubator. The medium was changed every 2-3 days. Cells were cultured in T-75 Falcon Tissue Culture Treated Flasks (Fisher Scientific, catalog #353112) until 75% confluent and split using TrypLE Express (TFS, catalog #12604013) for 5 min in the 37°C incubator. Cells were re-suspended in RPMI supplemented with 10% FBS and 1X penicillin-streptomycin and counted with a Countess Automated Cell Counter (TFS) using Countess™ Cell Counting Chamber Slides (TFS, catalog #C10228).

### Cell proliferation assay

A549 and H1975 cells were seeded at 1000 cells per well in a 6-well CytoOne tissue culture plate (USA Scientific, catalog #CC7682-7506) and cultured in full RPMI medium for 24 hours. After 24 hours, the medium was changed to mock treatment (full RPMI media), RPMI medium containing 10 µM H84T, or 10 µM of the largely inactive BanLec mutant D133G. The cells were cultured at 37°C in a 5% CO2 cell culture incubator for up to 5 days. On day 0, 3, 4 or 5, the cells were trypsonized with TrypLE Express, collected, and centrifuged at 1500 rpm for 5 mins. The cells were resuspended in full RPMI media and counted with a Countess Automated Cell Counter using Countess™ Cell Counting Chamber Slides. Cell lines were counted on different days due to differing rates of cell growth.

### 3D Culture

Cells were grown in 2% growth factor reduced Matrigel following the protocol previously described by Pal and Kleer (24). Briefly, precooled Lab-Tek 8-well glass chamber slides (TFS, catalog #177402) were coated with 40 µL of GFR Matrigel (BD Biosciences, catalog #354230) and then incubated at 37°C in a 5% CO_2_ cell culture incubator for 15 minutes, or until solidified. The cells were counted and diluted to 12,500 cells/mL in complete media of the respective cell line, GFR Matrigel was added to a working concentration of 2%, and 400 µL were gently added to each Matrigel pre-coated chamber. Phase contrast images were acquired at day 4-8 at 5X or 10X resolution on a Leica inverted microscope.

### Colocalization Assay

The NSCLC cell lines A549 and H1975 were seeded at 30,000 cells per well in a Millicell EZ slide 8-well glass and cultured in full RPMI for 24 hours. After 24 hours, the media was changed to mock treatment (full RPMI), RPMI containing 10 µM H84T or 10 µM D133G. The cells were treated for 30 minutes, 2 hours, and 4 hours. After removal of the treatment media, the cells were rinsed with full media, and washed two times with phosphate buffered saline (PBS, TFS, catalog #10010023). The cells were permeabilized with 0.5% Triton X-100 for 10 mins at 4°C and then washed three times with 100 mM glycine for 15 minutes each on a rocking platform at room temperature. The cells were blocked with 10% goat serum in immunofluorescence buffer (IF buffer, 1X PBS containing 0.1% bovine serum albumin, 0.2% Triton X-100, and 0.05% Tween-20) for 1 hour at room temperature followed by primary antibody incubation with anti-LAMP1 antibody, anti-EGFR antibody, or anti-His antibody for 1 hour at 4°C, followed by washing with IF buffer, 3 times for 15 minutes each, on a rocking platform at room temperature, and then incubation with secondary antibody (goat anti-mouse, goat anti-rabbit, or goat anti-rat antibodies) for 45 minutes at room temperature on a rocking platform. The cells were then washed 3 times with 1X PBS for 10 minutes each at room temperature. The slides were mounted using ProLong™ Gold AntiFade Mountant with DAPI (Invitrogen, catalog #P36935), and images were acquired using a Nikon A1 confocal microscope.

### Immunoblot Analysis

A549 cells were seeded on 10 mm cell culture plates and cultured in full RPMI for 24 hours. After 24 hours, the media was changed to mock treatment (full RPMI), RPMI containing 10 µM H84T or 10 µM D133G. The cells were treated for 72, 96, and 120 hours, washed twice with ice cold 1X PBS, and then collected with a cell scraper. The cells were lysed with lysis buffer that contained 0.1 M HEPES, 0.5 M NaCl, 0.5 M EDT, 0.25 M EGTA, 50% glycerol, 0.001% NP40, 0.001% SDS supplemented with complete Protease Inhibitor Cocktail (Roche, catalog # 11697498001) and protease inhibitor. Protein lysates, 33 µg per sample, were separated in Bio-Rad 4-20% Mini-PROTEAN® TGX gels (Bio-Rad, catalog #4561094). Proteins were transferred to 0.45 µm pore size polyvinylidene difluoride membranes (PVDF) (Millipore, catalog # IPVH00010), incubated with rabbit anti-LC3B antibody (1:1000, Cell Signaling Technology (CST), catalog #3868), rabbit anti-EGFR antibody (1:1000, CST, catalog # 54359), rabbit anti-Phospho EGFR antibody (1:1000, CST, catalog #4407), rabbit anti-AKT antibody (1:1000, CST, catalog #9272), or rabbit anti-Phospho AKT antibody (1:1000, CST, catalog #4060) overnight at 4°C, and then incubated with goat anti-rabbit secondary antibody (1:5000). As a loading control, the PVDF membrane was incubated with mouse anti-GAPDH antibody (1:10000, Santa Cruz, catalog #166545), and then incubated with horse anti-mouse secondary antibody (1:10000, CST, catalog #7076S).

### Apoptosis Assay

The NSCLC cell line A549 was seeded on 10 mm cell culture plates and cultured in full RPMI for 24 hours. After 24 hours, the media was changed to mock treatment (full RPMI), RPMI containing 2 µM Staurosporine, 10 µM H84T, or 10 µM D133G. The cells were treated with Staurosporine for 4 and 24 hours, while the H84T and D133G cells were treated for 24, 48, 72, 96, and 120 hours. The cells were washed twice with ice cold 1X PBS, collected with a cell scraper, and lysed with lysis buffer that contained 0.1 M HEPES, 0.5 M NaCl, 0.5 M EDTA, 0.25 M EGTA, 50% glycerol, 0.001% NP40, 0.001% SDS supplemented with cOmplete Protease Inhibitor Cocktail and protease inhibitor. Protein lysates, 33 µg per sample, were separated in Bio-Rad 4-20% Mini-PROTEAN® TGX gels. Proteins were transferred to 0.45 µm pore size polyvinylidene difluoride membranes (PVDF), incubated with rabbit anti-PARP antibody (1:1000, CST, catalog #9542), rabbit anti-C-PARP antibody (1:1000, CST, catalog #5624), rabbit anti-Caspase 3 antibody (1:500, CST, catalog #9664), or rabbit anti-Caspase 3 antibody (1:1000, CST, catalog #9662) overnight at 4°C, and then incubated with goat anti-rabbit secondary antibody (1:5000). As a loading control, the PVDF membrane was incubated with mouse anti-GAPDH antibody (1:10000), and then incubated with horse anti-mouse secondary antibody.

### Senescence Assay

The NSCLC cancer cell line A549 was seeded on a Millicell EZ slide 8-well glass and incubated for 24 hours. After 24 hours, the medium was changed to mock treatment (full RPMI media), RPMI medium containing 10 µM H84T or 10 µM D133G. The cells were cultured at 37°C in a 5% CO2 cell culture incubator for up to 14 days. The etoposide samples were incubated with full RPMI. Five days prior to ß-Galactosidase staining the samples were treated with 12.5 µM etoposide for 24 hours and allowed to recover for 4 days prior to ß-Galactosidase staining. For ß-Galactosidase staining the sample media were removed and the cells were washed once with PBS, and then fixed with 1X Fixative solution for 15 minutes at room temperature. The wells were then washed twice with 1X PBS and incubated with 1X ß-Galactosidase staining overnight in a dry incubator at 37°C. After 24 hours, the ß-Galactosidase stain was removed and replaced with 70% glycerol for storage. The slides were mounted with ProLong Gold AntiFade Mountant with DAPI, and images were acquired with a Nikon Eclipse E800 Upright Fluorescence microscope.

### Tissue Microarray

Formalin-fixed, paraffin-embedded tissue blocks (FFPE) of 136 cases of primary pulmonary cancer (76 adenocarcinoma and 60 squamous cell cancers) were obtained from the files of the Department of Pathology, University of Michigan Medical Center, Ann Arbor, MI. The University of Michigan Institutional Review Board provided a waiver of informed consent to obtain these samples. After pathological review, a tissue microarray (TMA) was constructed from the most representative area using the methodology of Nocito et al (25). Each case was represented by either two 1 mm diameter cores or three 0.7 mm diameter cores, obtained from the most representative, non-necrotic area of the tumor.

### Immunohistochemistry and scoring

Immunohistochemical staining was performed on the DAKO Autostainer (DAKO) using Envision+ and diaminobenzadine (DAB) as the chromogen. De-paraffinized sequential sections were labeled with either anti-EGFR (1:300, Rabbit monoclonal clone EP38Y, AbCam catalog #Ab52894) for 60 minutes at ambient temperature after incubation of the section with Background Sniper (BioCare Medical) for 30 minutes at ambient temperature or BanLec H84T as previously described (26). Microwave epitope retrieval in 10 mM sodium citrate buffer pH=6 was used prior to staining. Appropriate negative (no primary antibody) and positive controls (skin (EGFR) or NCI-H23 cell line pellet (BanLec H84T)) were stained in parallel with each set of slides studied.

Digital images were generated using an Aperio AT2 scanner (Leica Biosystems Imaging) at 20X magnification, with a resolution of 0.5 um per pixel. The scanner uses a 20x / 0.75 NA objective and an LED light source. Images were then analyzed using QuPath (27). Tumor associated immunoreactivity was assessed using the criteria of Harvey et al (28).

### Co-Expression of EGFR and BanLec H84T

Primary pulmonary adenocarcinoma TMA was serially stained with biotinylated BanLec H84T (red), EGFR (yellow) and pan-cytokeratin (green) and nuclei (blue) as previously described by McKenna et al (26). Between each staining step, the slide was subjected to heat induced epitope retrieval to remove the antibody/secondary antibody complex. The TMA was scanned using a Polaris fluorescence scanner. Separate images were analyzed using ImageJ to determine colocalization as previously described (26).

## Data availability

All data are contained within the manuscript.

## Results

### H84T BanLec inhibits the growth and proliferation of NSCLC cells

Cancers such as NSCLC present high mannose on their surfaces, making them potential targets for anti-glycan therapy (29). As H84T specifically attacks high mannose, is very well tolerated in mouse models even with repeated dosing (anti-BanLec antibodies develop but do not neutralize the molecule or harm the animals), goes directly to the lung when administered systemically, and has a 35-hour half-life in blood and lung, we hypothesized that H84T could be an effective therapy for NSCLC (17, 18).

We therefore tested the ability of H84T to inhibit the growth of lung cancer cell lines in tissue culture, and found that the inhibitory effects are striking, as we first observed in 3-dimensional systems. The spheroid-like colonies of lung cancer cell lines that grow in 3D culture have been reported as a physiologically relevant tumor-mimicking tool and have increasingly been used for anti-cancer drug screening purposes (30). The colony forming ability of the lung cancer cell lines A549 (wild-type EGFR, Figure 1A), H1975 (which harbors the EGFR mutations L858R, targeted by EGFR TKIs, and T790M, which confers resistance to first and second generation TKIs but is susceptible to osimertinib, Figure 1B), and HCC827 (harbors EGFR exon 19 deletion, also targeted by EGFR TKIs, Figure 1C) was markedly inhibited by H84T, but not by the D133G (non-mannose binding) BanLec (20). To further confirm the activity of H84T against NSCLC cells, and to use a system that more readily lends itself to the study of signal transduction and cell death pathways, we examined whether H84T is effective against NSCLC cells in 2-dimensional, conventional tissue culture, and found that here too it is highly active (Figure 2). Proliferation of both A549 (Figure 2A) and H1975 (Figure 2B) was inhibited by H84T, an effect that became more pronounced after 4 and 5 days of treatment.

**Figure 1.**
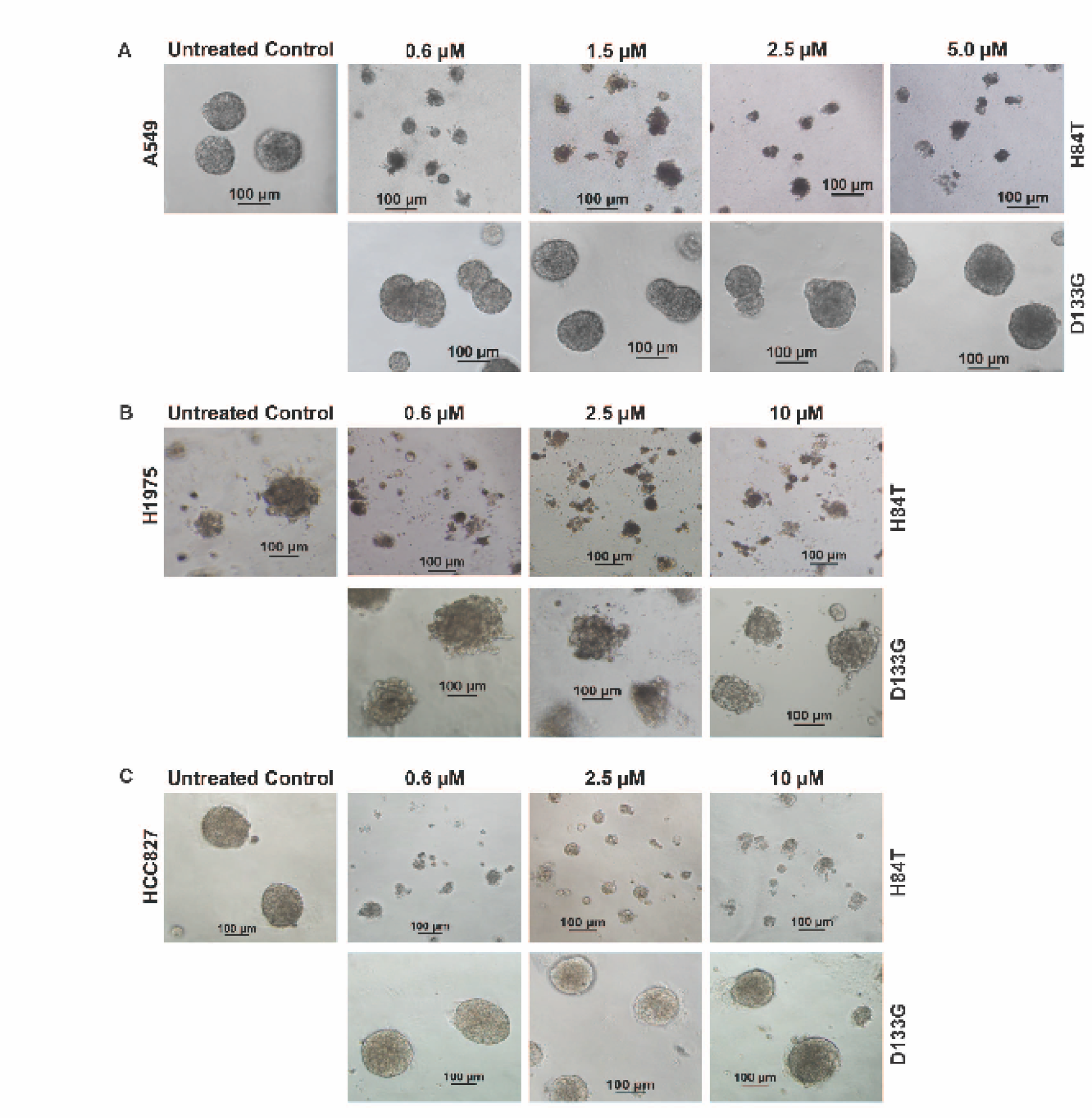
Treatment of lung cancer cell lines with H84T inhibits colony growth in 3-dimensional cultures. (**A**) A549, NSCLC cell line expressing wild-type EGFR, (**B**) H1975, NSCLC cell line with an activating mutation of EGFR that confers resistance to first- and second-generation TKI, and (**C**) HCC827, NSCLC cell line with an activating EGFR deletion, were cultured in 3-dimensions on 2% growth factor reduced Matrigel in the absence (control) or presence of varying doses of purified H84T or D133G. The viability of the colonies was evaluated with phase-contrast microscopy 2-6 days after treatment. At the lowest dose administered (0.6 µM), colony growth was almost completely suppressed in each of the cell lines in the presence of H84T, demonstrating EGFR mutation-independent activity of H84T. D133G, a single residue variant of BanLec that largely lacks mannose binding, has no cytotoxic activity, even at the highest dose tested.

**Figure 2.**
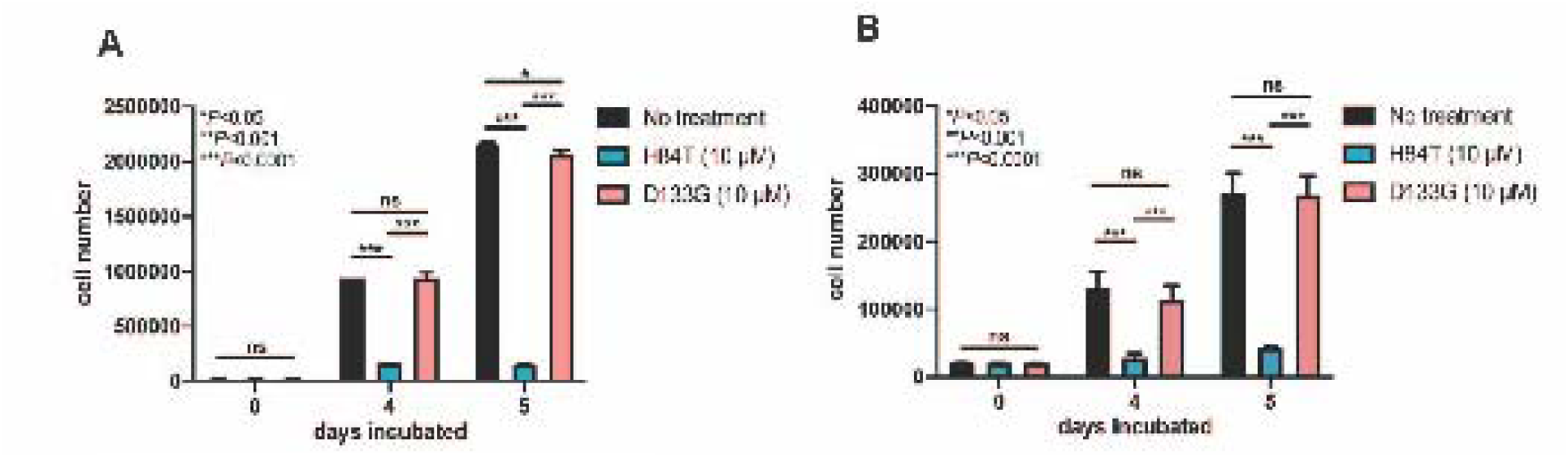
Treatment of lung cancer cell lines with H84T inhibits cell proliferation. (**A**) A549 and (**B**) H1975 lung cancer cells were cultured in the absence (control) or presence of 10 µM H84T or D133G BanLec. Cell proliferation of A549 and H1975 cells was evaluated by cell counting on days 0, 4 and 5. Treatment NSCLC cells with H84T significantly reduced cell proliferation after 4 and 5 days of treatment in both A549 (p<0.0001 for both timepoints) and H1975 cells (p<0.0001 for both timepoints).

### Treatment of NSCLC cells with H84T BanLec leads to the degradation of EGFR and inhibition of AKT signaling

EGFR is a receptor that displays high mannose specifically in transformed cells and is vital to the growth of a number of different tumors, including NSCLC (3, 4). We investigated the mechanistic basis for the inhibition of NSCLC cell proliferation observed in Figures 1 and 2. Adding H84T to the NSCLC cell line A549 led to degradation of EGFR (Figure 3). The inhibition of lung cancer growth observed in Figures 1 and 2 thus correlated with the degradation of EGFR. Furthermore, treatment of NSCLC cells with H84T led to the subsequent downregulation of AKT expression levels. We found that the treatment with H84T, but not the inactive (non-mannose binding) D133G mutant of BanLec, led to the degradation of EGFR and inhibition of AKT signaling pathways in NSCLC cells at all timepoints (Figure 3). Figure 3 also shows changes in LC3B indicating that autophagy is the likely mechanism of cell death; this is further discussed below.

**Figure 3.**
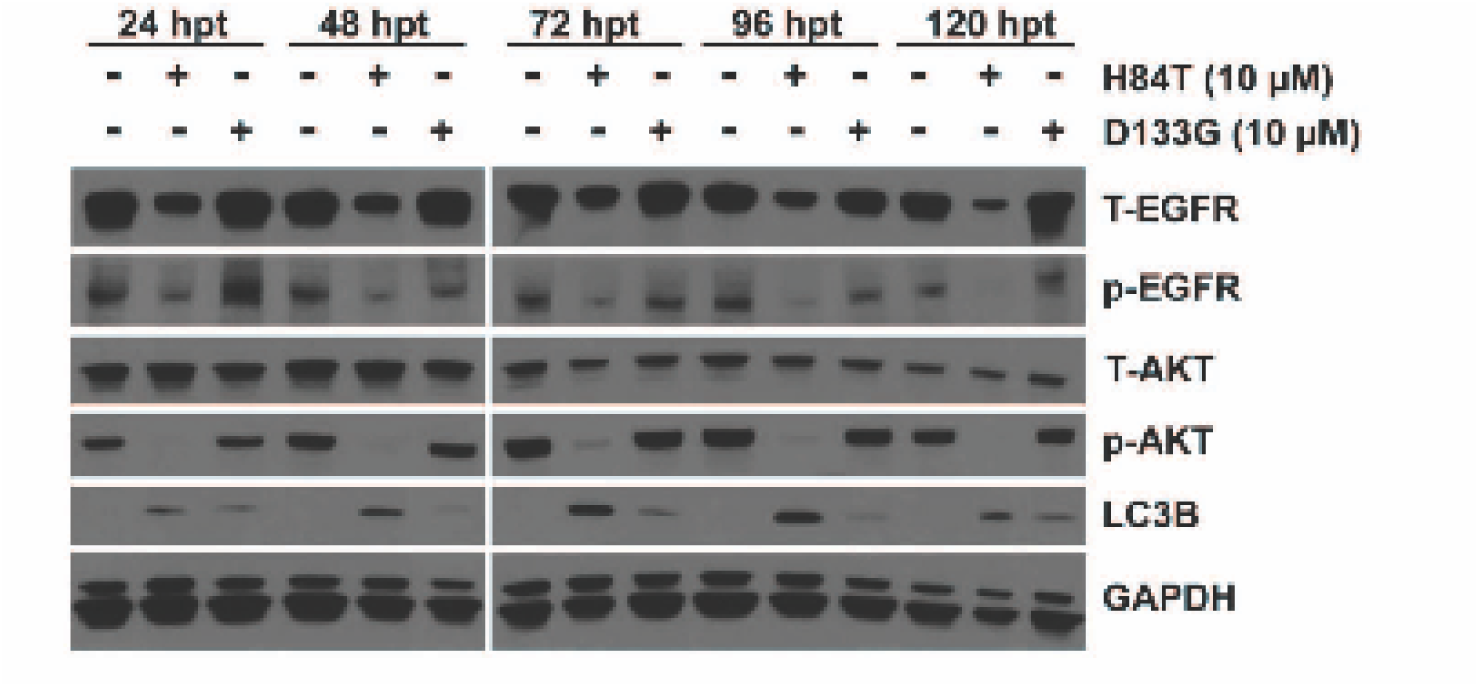
Treatment of NSCLC cells with H84T BanLec leads to the degradation of EGFR and inhibition of AKT signaling and induces autophagy. A549 cells were cultured in the absence (control) or presence of 10 uM H84T or D133G for 24, 48, 72, 96 or 120 hours. At all timepoints, treatment of A549 cells with H84T resulted in degradation of both phospho (p)-EGFR and total (T)-EGFR, and downregulation of p-AKT. Additionally, treatment of A549 cells with H84T (and to a lesser degree D133G) resulted in increased levels of LC3B, a known autophagosome marker.

### H84T, but not D133G, binds to EGFR expressing cells, is internalized, and colocalized in the lysosomal compartment

Importantly, H84T, but not D133G, bound to EGFR on the cell surface of the NSCLC cell line A549 (Figure 4), and led to its internalization into an endocytic compartment apparently leading to lysosomal degradation (Figure 5), a mechanism quite distinct from the EGFR TKIs that are currently in clinical use. Colocalization of H84T with EGFR was present in a time-dependent manner. Treatment of A549 cells with H84T, but not D133G, showed colocalization of H84T with EGFR in the lysosomal compartment, as confirmed by the colocalized staining of LAMP1, a known lysosomal marker. After 4 hours of treatment, there was robust co-staining of H84T and EGFR with LAMP1. D133G did not bind to the cell surface and did not affect EGFR or LAMP localization (Figure 5).

**Figure 4.**
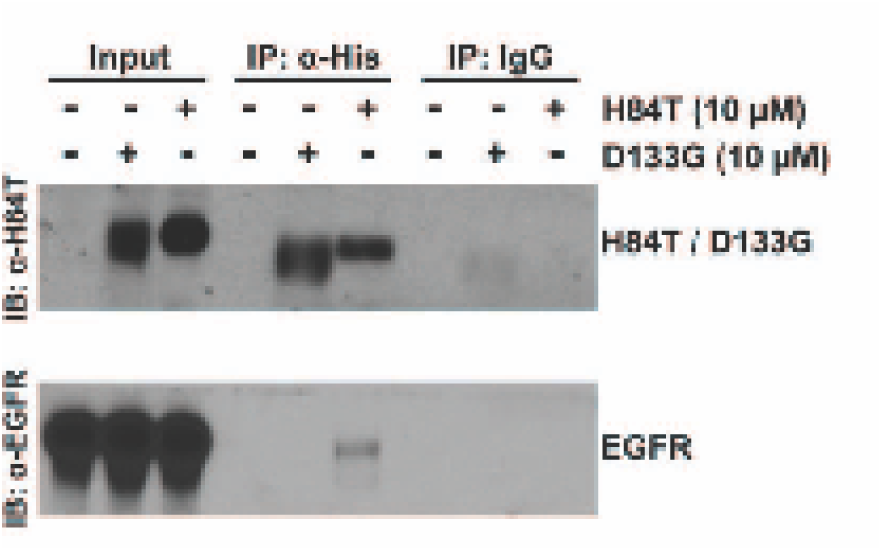
H84T binds to EGFR. A549 cells were treated with 10 uM H84T or D133G. Cell extracts were prepared and used for immunoprecipitation of H84T or D133G BanLec with an anti-His antibody or, as a control, a non-specific IgG antibody. Analysis of the extracts (input) demonstrated that EGFR and BanLec were present. However, immunoprecipitation of BanLec resulted in co-immunoprecipitation of EGFR only in extracts containing H84T but not D133G. This demonstrates that H84T, but not D133G, binds to EGFR.

**Figure 5.**
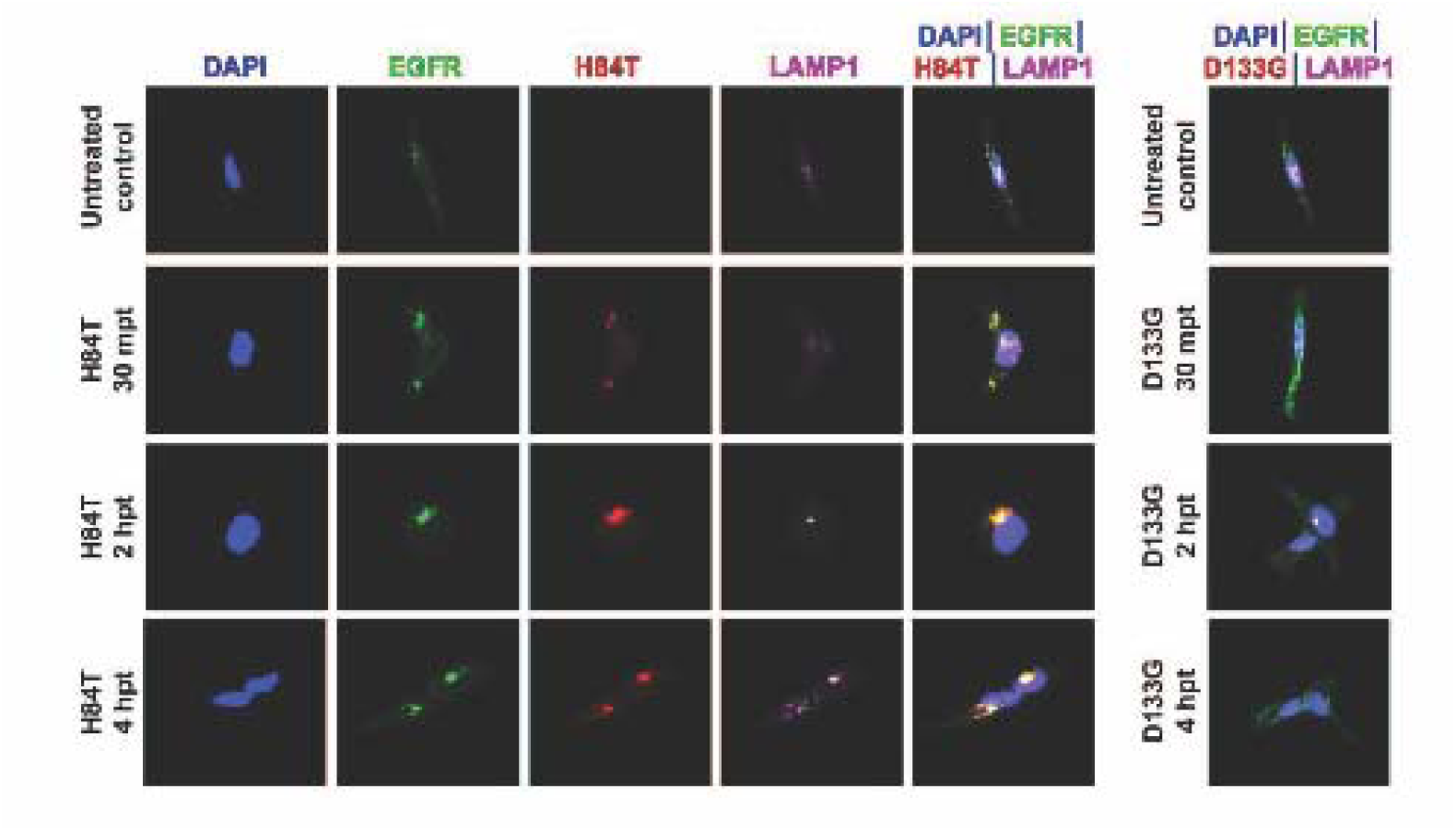
H84T but not D133G binds to EGFR expressing cells, is internalized, and colocalizes in the lysosomal compartment. A549 cells were treated with 10 uM H84T or 10 uM D133G and fixed for immuno-cytochemistry at 30 minutes, 2 hours, and 4 hours post treatment. At 30 minutes, plasma membrane staining of H84T (red) and EGFR staining (green) began to colocalize (yellow-orange) with LAMP1 (fuchsia) in the endosomal-lysosomal department. At 2 hours, an increase in colocalized staining of EGFR and H84T with the LAMP1 lysosomal marker (yellow signal) was observed intracellularly. At 4 hours, a robust staining of EGFR and H84T signal was observed to be colocalized in the lysosome. D133G did not bind to the cell surface and did not alter EGFR or LAMP1 localization.

### Treatment of BaF3/EGFR cells with H84T leads to inhibition of cell growth

The Ba/F3 cell line is often used as a model system for assessing the effects of kinase oncogenes and their sensitivities to inhibitors (31). Normal Ba/F3 cells do not express EGFR and are refractory to EGFR-inhibitors. However, EGFR expressing BA/F3cells (BaF3/EGFR) demonstrate potent sensitivity to EGFR inhibition. To investigate the sensitivity of EGFR-addicted cells to H84T and its inactive mutant D133G, we assessed growth inhibition using the CellTiter-Blue® Cell Viability Assay. Indeed, H84T, but not D133G, was cytotoxic to BaF3/EGFR cells (Supporting Figure S1). Furthermore, H84T but not D133G was cytotoxic to BAF3-L858R/C797S-EGFR (osimertinib resistant) and BaF3/EGFR-L858R/T790M/C797S-EGFR (osimertinib resistant) cells (Supporting Figure S1).

### H84T BanLec treatment induces autophagy in NSCLC cells

We next examined the mode of death for the H84T treated A549 cells. H84T treatment resulted in increased expression of LC3B, an autophagosome marker that reflects autophagic activity (Figure 3). Interestingly, D133G, which has no antiviral activity and does not inhibit NSCLC growth, does induce a certain degree of autophagy, perhaps due to non-carbohydrate binding effects. Furthermore, treatment of NSCLC cells with H84T BanLec did not induce apoptosis (Supporting Figure S2) or senescence (Supporting Figure S3), confirming that autophagy is the major mode of NSCLC death in response to H84T treatment of NSCLC. Interestingly, D133G did induce some late apoptosis (just as it did autophagy), despite its inability to inhibit cell growth.

### H84T directly interacts with diverse NSCLC samples from patients

We hypothesized that because EGFR on NSCLC cells is covered with high mannose, whereas EGFR on healthy cells does not bear high mannose regardless of expression level, H84T would be specific for tumors from patients with NSCLC and not for healthy lung tissue. Indeed, we have previously found minimal to no interaction between H84T and healthy human, primate, and rat tissues (18, 21, 22). We thus performed immunohistochemical analysis of NSCLC and healthy lung tissue on a TMA with >100 tissue specimens and found that indeed H84T reacts strongly with both types of NSCLC (adenocarcinoma and squamous cell carcinoma) (Figure 6A and B), whereas healthy lung tissue only minimally reacts with H84T (Figure 6C). Interestingly, H84T does recognize macrophages, as would be predicted because these are some of the few healthy human cells that present high mannose. We have previously shown that the interaction of H84T with hematopoietic cells is not functionally relevant (23).

**Figure 6.**
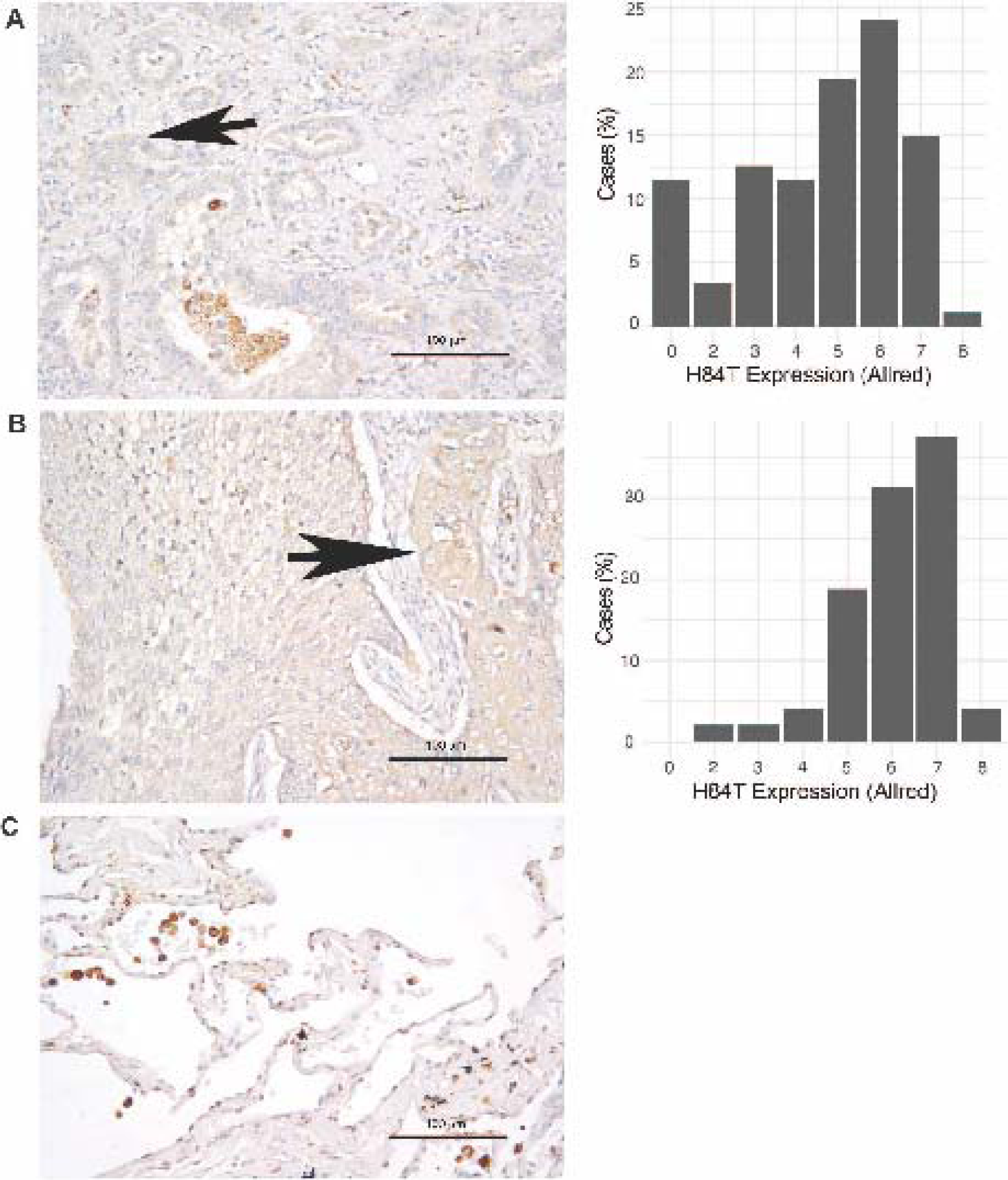
H84T binds to human NSCLC, but not to healthy lung tissue. Sections of pulmonary adenocarcinoma and squamous cell carcinoma TMAs were stained with H84T BanLec. Stained slides were digitized and automatically scored using QuPath. Allred Scores were binned and tabulated. **(A)** Immunohistochemical staining using biotinylated H84T to probe primary pulmonary adenocarcinoma tissue, highlighted by arrow (L) and Allred score (R) **(B)** H84T staining of primary pulmonary squamous cell carcinoma tissue, highlighted by arrow (L) and Allred score (R) **(C)** Healthy pulmonary tissue exhibits minimal staining by H84T; only alveolar macrophages stained positive. It is well known that alveolar macrophages express high mannose.

H84T shows strong colocalization with EGFR in NSCLC cell lines, leading to the destruction of this receptor (Figure 4 and Supporting Figure S1). We therefore used the TMAs to examine whether such colocalization is seen in actual human tumor samples. Indeed, we found that H84T has an extremely high degree of colocalization with EGFR in human primary lung adenocarcinoma tissue (Spearman ρ=0.98; Figure 7), confirming that the effects we see in NSCLC cell lines hold true for human lung cancer specimens as well.

**Figure 7.**
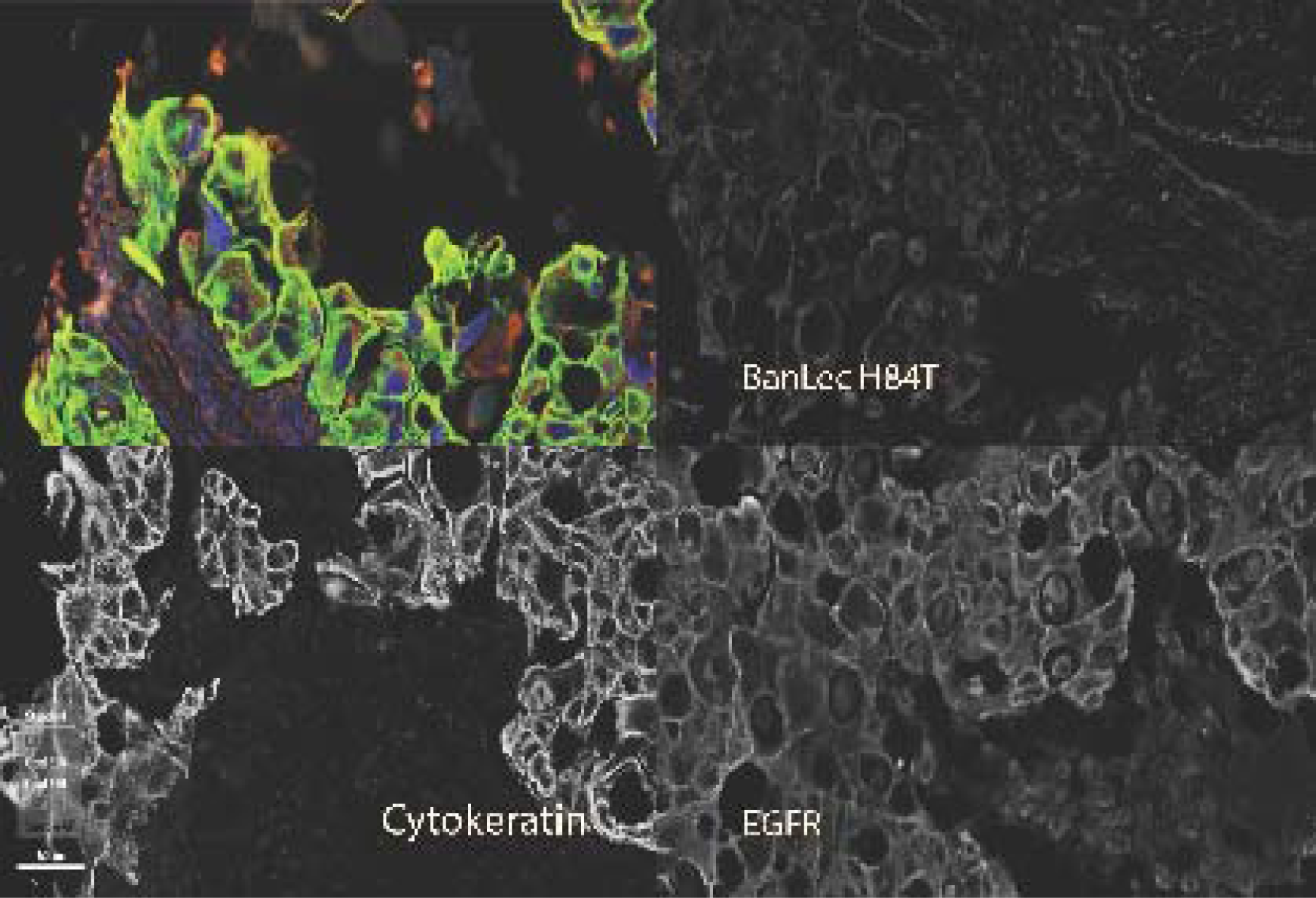
H84T colocalizes with EGFR in human NSCLC tissue. A primary pulmonary adenocarcinoma tissue microarray (TMA) was serially stained for biotinylated H84T BanLec (red), EGFR (yellow), pan-cytokeratin (green), and nuclei (blue). Between each staining step, the slide was subjected to heat induced epitope retrieval to remove the antibody/secondary antibody complex. The TMA was scanned using a Polaris fluorescence scanner. A representative core is show in this figure (original magnification 200x), demonstrating colocalization of the lectin with both EGFR and pan-cytokeratin. Separate images were analyzed using ImageJ to determine colocalization with a Spearman’s rank correlation value (ρ) of 0.98.

## Discussion

While great strides have been made in the treatment of NSCLC with the advent of EGFR TKIs, resistance invariably develops and survival rates remain low; additionally, these therapies can only be used in the minority of patients with susceptible driver EGFR mutations. As EGFR is overexpressed in the majority of NSCLC and is heavily implicated in pathogenesis, it remains a promising target for an agent that can overcome or circumvent the challenges faced by TKIs (32).

Abnormal glycoproteins on the surface of cancer cells are promising therapeutic targets, as these glycoproteins are often essentially unique to malignant cells, allowing for precision targeting of the tumor and limiting the potential for off-target effects. However, advancement of glycan-targeting cancer therapies has been hampered by lack of glycan-binding agents with sufficient specificity for the target (12). Lectins have evolved precisely for this purpose and are able to bind glycans with high specificity. Lectins are already used to detect abnormal glycans in cancer diagnostics and prognostics, and there have been recent developments in the use of lectins as cancer therapeutics as well, including using H84T to target pancreatic cancer with CAR-T therapy (12, 13, 23, 33–35).

H84T is a lectin derived from bananas previously engineered by our group to separate its antiviral activity from its mitogenicity. H84T exerts its antiviral activity by binding high mannose found on viral surfaces (20). As EGFR in NSCLC, like many cancer-associated proteins, expresses high mannose – which critically is not expressed on most healthy cells – we hypothesized that H84T could be an effective treatment for NSCLC (13, 14).

We have now shown that H84T effectively binds to, and inhibits the growth of, NSCLC cells through a mechanism distinct from the TKIs currently in clinical use. Treatment with H84T markedly inhibited the growth of multiple NSCLC cell lines. Importantly, H84T was able to significantly reduce the proliferation of NSCLC cells that do not express EGFR driver mutations, and therefore would not be targeted by TKIs. This is likely because high mannose expression on the surface of malignant EGFR seems to be independent of mutation status and is present on tumor-associated wild-type EGFR as well (13, 36). This is a meaningful finding as the majority of patients do not express these mutations and are not candidates for TKIs; H84T is a potential treatment for these patients. Notably, H84T was also able to inhibit the growth of osimertinib-resistant NSCLC cell lines. As noted above, patients invariably develop resistance to TKIs, thus a treatment that is not susceptible to these resistance pathways or could be used after resistance develops would also be a significant development. We should also note that there are surface receptors other than EGFR that bear high mannose in NSCLC (and other malignancies) and so, while EGFR is a major target of H84T, it is unlikely to be the only one. Indeed, insulin-like growth factor 1 receptor, another receptor known to contribute to NSCLC pathogenesis, was also recently shown to be targeted by a high mannose-binding lectibody (13).

H84T exerts its effect on NSCLC growth by inducing the lysosomal degradation of EGFR. We have shown that H84T binds to EGFR on the surface of NSCLC and subsequently leads to internalization and co-localization of EGFR with the lysosome, and ultimately to EGFR degradation and inhibition of downstream signaling pathways. This is a mechanism distinct from EGFR TKIs, which act by either reversibly binding the EGFR kinase domain and therefore interrupting downstream signaling (first-generation TKIs), irreversibly blocking EGFR transphosphorylation (second-generation TKIs), or irreversibly binding specific mutant forms of EGFR (third-generation TKIs) (6). Mechanisms of resistance, especially to third generation TKIs, are complex and incompletely understood, but largely involve either mutations in EGFR itself or in downstream pathways (6, 10). Inhibition through lysosomal degradation is unlikely to be subject to the same resistance mechanisms. Lysosomal degradation of EGFR has been reported as a mechanism of NSCLC suppression by both endogenous signaling pathways and putative therapeutics (37–39). Indeed, restoration of lysosomal degradation has been reported to overcome EGFR TKI resistance both in vitro and in mouse models of NSCLC, and represents a promising mechanism for EGFR-directed therapeutics (40, 41).

We also found that H84T induces autophagy in NSCLC cells. Autophagy, the homeostatic process by which cells degrade and recycle proteins, damaged organelles, and other components via lysosomal degradation, has a complex relationship with cancer. In some contexts, autophagy promotes cancer cell survival and growth, and in others autophagy works to limit cancer establishment and progression (42, 43). EGFR itself is thought to be a critical factor in whether autophagy is protective or oncogenic (43). Similarly, what role autophagy might play in EGFR TKI pathways is not clear. Some studies have reported that increased autophagy mediates EGFR TKI resistance, and that inhibition of autophagy potentiates the effects of TKIs. However, autophagy-mediated cell death also seems to be a key mechanism by which EGFR TKIs exert their effects (44–49). The present study demonstrates that H84T induces NSCLC cell death through autophagy. We have additionally shown that H84T does not induce senescence or apoptosis in NSCLC cells, further indicating that autophagy is likely responsible for H84T-induced growth inhibition.

As H84T colocalizes with EGFR in the lysosome, leading to EGFR degradation, it seems likely that this event is upstream of H84T-induced autophagy. Akt, which is a critical downstream effector of EGFR as part of the PI3K/Akt/mTORC1 pathway, is also downregulated in H84T-treated cells. The PI3K/Akt/mTORC1 pathway is one of the mechanisms by which EGFR regulates autophagy in cancer cells, but as noted above, the effects are complex and difficult to predict (43). While Akt is typically considered to be a negative regulator of autophagy, this pathway has been shown to both induce and inhibit cancer-related autophagy in different circumstances; both effects have been shown in NSCLC cells specifically (44, 50). Given that H84T both decreased p-Akt and induced autophagy in the absence of another apparent mechanism of cell death, it appears likely that inhibition of Akt is associated with increased autophagy, a mechanism that has also been reported with EGFR TKIs (44).

Significantly, H84T recognizes with high affinity the large majority of NSCLC specimens obtained from patients. Such recognition is seen with both adenocarcinoma and squamous cell carcinoma of the lung, suggesting the possibility of personalized but broad clinical utility. We further found that H84T is specific for NSCLC tissue and does not bind healthy lung tissue. This observation is expected as H84T is highly specific for high mannose, which is not found on most healthy cells, and is in line with the fact that we have safely treated hundreds of mice, hamsters, and rats in our antiviral studies of H84T (18, 21, 22). This specificity is critical, as many of the adverse effects of EGFR TKIs are mediated by binding EGFR on healthy tissues as well as on cancer cells (though these adverse effects are moderated in third generation TKIs, which are more specific for malignant EGFR) (6). As high mannose is expressed on many cancer cell surfaces and not only on NSCLC-associated EGFR, it is notable that we have found that H84T is able to exert similar effects in reducing cell proliferation and survival across many cancer cell lines, including breast cancer, pancreatic cancer, and hepatocellular carcinoma, without binding to normal, healthy tissues (data not shown). Importantly, the inactive banana lectin D133G, a single residue variant of H84T that does not bind high mannose, does not bind EGFR and has no effect on inhibiting the proliferation of NSCLC cells, although it does induce a small degree of autophagy and even apoptosis. Why we see these effects is unclear; perhaps D133G BanLec maintains some small degree of lectin activity or affects autophagy/apoptosis through non-sugar binding mechanisms that are not sufficient to block cell growth.

In conclusion, this study shows that H84T, a molecularly modified lectin, binds high mannose on the surface of NSCLC cells, induces lysosomal degradation of EGFR, and leads to cancer cell death through autophagy. This effect is independent of EGFR mutation status (including mutations that confer TKI resistance), and therefore has the potential to be effective for the majority of NSCLC. These effects are specific to malignant cells as H84T does not bind normal, healthy tissue. H84T, which is well-tolerated in animal models of viral infection, is a promising new potential therapeutic for a disease where novel therapies are urgently needed (17–22).

## Supporting Information

This article contains supporting information.

## Funding and additional information

ZR was supported by NIH grant T32HL007749. MKN was supported by NIH grant R01CA248310. DMM was supported by NIH grant R01AI175124, a grant from the Forbes Institute of the Rogel Cancer Center at the University of Michigan, and by a Frankel Innovation Award.

## Dedication

This paper is dedicated to the memory of Dr. Robert A. Ketroser.

