## Supplementary figures and images for "A molecularly engineered lectin destroys EGFR and inhibits the growth of non-small cell lung cancer"

### Supplementary Figures

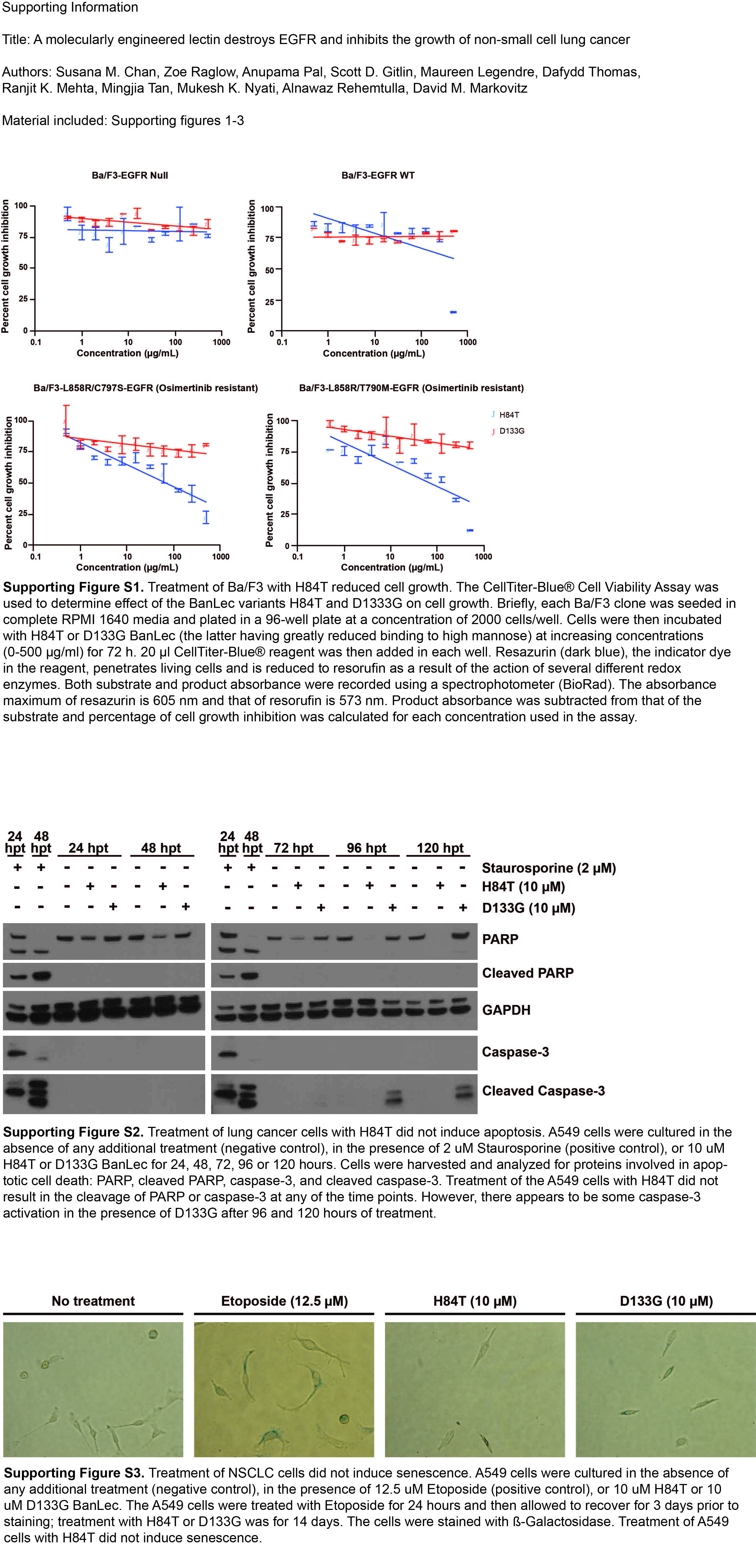
